# Estimating the overall fraction of phenotypic variance attributed to high-dimensional predictors measured with error

**DOI:** 10.1101/2022.02.25.482026

**Authors:** Soutrik Mandal, Do Hyun Kim, Xing Hua, Shilan Li, Jianxin Shi

## Abstract

In prospective genomic studies (e.g., DNA methylation, metagenomics, and transcriptomics), it is crucial to estimate the overall fraction of phenotypic variance (OFPV) attributed to the high-dimensional genomic variables, a concept similar to heritability analyses in genome-wide association studies (GWAS). Unlike genetic variants in GWAS, these genomic variables are typically measured with error due to technical limitation and temporal instability. While the existing methods developed for GWAS can be used, ignoring measurement error may severely underestimate OFPV and mislead the design of future studies. Assuming that measurement error variances are distributed similarly between causal and noncausal variables, we show that the asymptotic attenuation factor equals to the average intraclass correlation coefficients of all genomic variables, which can be estimated based on a pilot study with repeated measurements. We illustrate the method by estimating the contribution of microbiome taxa to body mass index and multiple allergy traits in the American Gut Project. Finally, we show that measurement error does not cause meaningful bias when estimating the correlation of effect sizes for two traits.

## 1. Introduction

Genome-wide association studies (GWAS) aim to identify genetic variants associated with a specific phenotype by examining millions of genetic variants across the genomes^1,2^. Since the first GWAS was successfully performed for age-related macular degeneration in 2005^3^, hundreds of associations for many diseases have been identified. These findings have improved our understanding of disease etiology and genetic risk prediction. While GWAS have been successful as a revolutionary method for genetic mapping of complex traits and diseases, the identified few variants can only explain a small fraction of heritability estimated based on twin or family studies. This so-called “missing heritability” problem puzzled scientists. By that time, the identified few variants could only explain a small fraction of heritability estimated based on twin or family studies. For example, the approximately 180 genetic variants identified in large-scale GWAS of human height published in 2010 explained about 10% of phenotype variance^4^, much less than the 80% heritability estimated based on twin- or family-based studies. This raised a question whether GWAS would be highly productive and provide important tools for genetic risk prediction eventually.

To answer this question, linear mixed models (LMM) were developed to estimate the overall fraction of phenotypic variation (OFPV), also called narrow sense heritability in genetics, explained by all common variants that are genotyped or imputed in a GWAS^5^. Here, OFPV corresponds to the variance component in LMMs. For human height, they estimated the heritability to be approximately 45%^5^, much higher than the 10% based on the 180 variants identified by that time^6^. This analysis provided compelling evidence that majority of causal variants were not detected because of insufficient power. For case-control studies, we can use this framework to estimate OFPV at the observational scale and then convert it to population-level OFPV by adjusting for ascertainment and disease prevalence in the target population^7^.

Recently, large-scale prospective studies with high-dimensional genomic variables have been performed or planned for various diseases. These prospective studies aim to identify genomic variables associated with disease risk to better understand disease etiology and to improve risk prediction. Examples of the genomic variables include but are not limited to microbiome taxa relative abundances, gene expression, microRNA expression, and DNA methylation. Typically, none or just a few genomic variables can be identified to be statistically significant even in a reasonably large-scale study, and the resulting predictive models have a poor performance. For example, a recent prospective peripheral blood DNA methylation study^8^ with 1,663 breast cancer cases and 1,885 controls at 365,145 CpG markers did not identify any significant association after multiple testing correction. In our recent study for never smokers from two cohorts – Shanghai Women’s Health Study and the Shanghai Men’s Health Study, we did not identify significant associations between oral microbiome and lung cancer risk^9^. In another oral microbiome study nested in cohorts, we identified a few significant associations for lung cancer risk^10^, but these associations only explained a small fraction of lung cancer risk. Similar to the puzzling “missing heritability” problem in GWAS, the lack of findings in these genomic studies may be caused by limited statistical power due to insufficient sample size or by the fact that the genomic variables do not contribute to the disease risk. Thus, it is informative to estimate OFPV, which helps decide the necessity and the scale of future studies.

While the statistical framework and software tools^5,11^ developed for GWAS can be readily used, there is important difference between these genomic variables and genetic variants in GWAS. Genotypes of common genetic variants remain unchanged over the whole life and can be accurately captured by genotyping arrays. However, genomic variables are typically measured with error, which comes from two sources: technical variability and temporal instability. Ignoring measurement error in genomic variables will cause a biased estimate of OFPV.

The problem of measurement error in independent variables, or errors-in-variables models, have been intensively investigated for regression analysis^12^, LMMs^13^ and generalized LMMs^14,15^. For a simple linear regression, measurement error in the independent variables attenuates the estimate toward zero; the attenuation factor or the reliability ratio^12^ equals to the intraclass correlation coefficient (ICC) of the variable. Treatment of measurement error in generalized linear models and other nonlinear models has been well discussed^16,17^. For LMMs or generalized LMMs, methods have been developed to correct for the bias in both fixed effects and variance components when there is measurement error in fixed effect variables^13-15,18^. When there is measurement error in both fixed and random effects, Cui and his colleagues developed moment estimators to consistently estimate both fixed and random effects^19^. While this framework is most relevant to our problem, it cannot be directly applied. The LMMs assume that all random effect variables are causal; however, signals are expected to be sparse in genomic studies^20^. As was pointed out by Jiang and his colleagues^21^, the LMMs^5^ used for estimating heritability are misspecified and statistical inference needs to be performed under the assumption of sparse signals. Moreover, the covariance structure of measurement error is assumed to be known^19^. In practice, this needs to be estimated using a dataset of limited sample size and the uncertainty in the estimate needs to be modeled when constructing confidence intervals.

In this manuscript, we aim to derive the asymptotic attenuation factor (AAF) for estimating OFPV when the high-dimensional genomic variables are measured with error. Because bias analysis is complicated based on the LMMs using restricted maximum likelihood (REML), we proceed with a recently developed pairwise regression that is similar to the Haseman-Elston regression^22,23^ in statistical genetics. We will show that AAF equals to the average ICC of all genomic variables under the assumption that the distributions of ICCs are identical between causal and non-causal variables. Thus, to derive a consistent estimator for OFPV, we need to perform a pilot study with repeated measurements to estimate the average ICC.

A parallel question is how to estimate the correlation of effect sizes between two traits, which was also motived in GWAS for analyzing genetic correlation between two traits^24,25^. We assume that the two traits are influenced by the same set of *K* genomic variables with effect sizes following a bivariate normal distribution with correlation *ρ*. Estimating *ρ* is crucial for understanding the degree of the shared etiology between the two traits. A joint study can be performed to achieve a high statistical power for detecting individual associations or risk prediction^26-29^ if multiple traits share the same etiology with a large value of *ρ* between primary and the secondary traits. Our analysis shows that measurement error has little impact for estimating *ρ* even if we ignore measurement error.

This manuscript is organized as follows. In Section 2, we derive AAF in the presence of independent and additive measurement error in genomic variables. In Section 3, we discuss estimating AAF using repeated measurements when ICCs are unknown. In Section 4, we investigate the impact of measurement error on estimating *ρ* for two traits. In Section 5, we present numerical studies. In Section 6, we present a data example estimating OFPV for body mass index (BMI) and multiple allergy traits explained by gut microbiome using the data from the American Gut Project. Limitations and future work are discussed in Section 7.

## 2. Estimating the overall fraction of phenotypic variance attributed to high-dimensional genomic variables

### 2.1 Notations and models

For subject *i* (*i* = 1, ⋯ *N*), let *y*_*i*_ denote the phenotypic value and (*x*_*i*1_, ⋯, *x*_*iM*_) denote the values of *M* independent genomic variables. Here, *y*_*i*_ and *x*_*im*_ have been normalized to have mean zero and unit variance. We consider an additive model *y*_*i*_ = ∑ _*m*∈*C*_ *β*_*m*_ *x*_*im*_ + *ε*_*i*_, where *C* denotes the set of *K* causal variables. In the narrow sense, 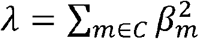 represents the OFPV explained by all *K* causal variables. Assuming a random effect model *β*_*m*_ ∼ *N*(0, *σ*^2^) *i*.*i*.*d*., then *λ* = *Kσ*^2^. One can show that *cor*(*y*_*i*_, *y*_*j*_) = *E* (*y*_*i*_*y*_*j*_) = *λg*_*ij*_, where

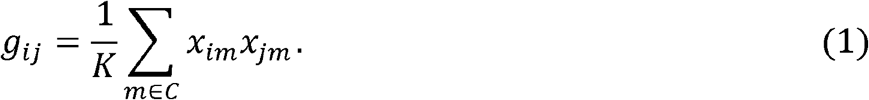

We define an *N* × *N* similarity matrix *G* with (*G*)_*ij*_ = *g*_*ij*_. Then, (*y*_1_, ⋯, *y*_*N*_) follows a multivariate normal distribution with covariance matrix *λG* + (1− *λ*) *I*.

Two types of methods have been proposed to estimate *λ*. The first method is the restricted maximum likelihood (REML) method. For GWAS, this method was implemented in a software package Genome-wide Complex Trait Analysis (GCTA)^5^ and has been used extensively. The second method regresses the empirical <u>P</u>henotypic <u>C</u>orrelations *y*_*i*_*y*_*j*_ onto the <u>G</u>enetic <u>C</u>orrelation (PCGC)^30^ *y*_*i*_*y*_*j*_ ∼ *λg*_*ij*_ for all *N* (*N* − 1)/2 pairs of subjects to obtain

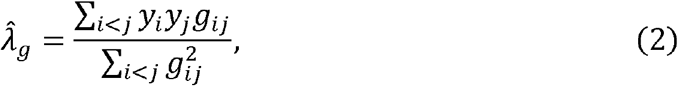

which is also called the Haseman-Elston regression in statistical genetics literature^22,23^. Existing studies showed that both methods are consistent^21,30^. We call 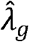 as the PCGC estimator. Note that the REML method is based on the full likelihood and provides a more efficient estimate than the PCGC method that uses only pairwise information. However, technical analysis of measurement error is easier for the PCGC estimator; thus, our investigation will be based on the PCGC estimator.

### 2.2 Estimating OFPV when the causal set *C* is known

We extend the PCGC method to the scenario when genomic variables are measured with error. For the genomic variable *m* and subject *i*, let *x*_*im*_ be the underlying unobserved true value, *e*_*im*_ be the measurement error and 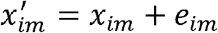 be the observed value of the error-prone variable.

We make three assumptions: (1) *e*_*im*_ and *x*_*im*_ are independent, (2) *e*_*im*_ and *e*_*in*_ are independent for any (*m, n*), and (3) 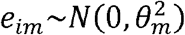. Define 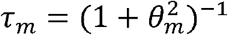, the ICC of the variable. Again, we normalize the observed value to have mean zero and unit variance

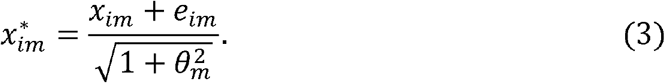

The error-prone version of *g*_*ij*_ in (3) is given by

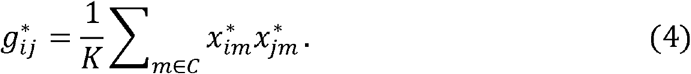

Let 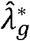 be the PCGC estimator based on 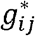 using formula (4). We will derive AAF for 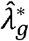.

#### Theorem 1.

*We assume that x*_*im*_, *e*_*im*_ (*i =*1, ⋯, *N*; *m* =1, ⋯, *M*) *are independent random variables with x*_*im*_ ∼ *N* (0,1) *and* 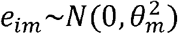. *Define* 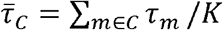 *be the average ICC for the causal genomic variables. Let* 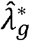 *be the PCGC estimator based on error-prone* 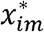 *defined in (3). As N, K* → ∞ *in a way that* 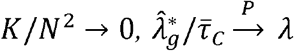.

Thus, 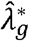 using error-prone predictors underestimates *λ* with 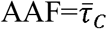. If we know 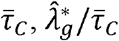 provides a consistent estimate of *λ*. As a special case, when all causal genomic variables have the same 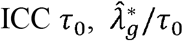 provides a consistent estimate of *λ*. Of note, the convergence requires *K*/*N*^2^ → 0 as *N, K* → ∞, i.e., the number of causal genomic variables does not increase too fast compared to the sample size *N*.

### 2.3 Genomic variables are measured with error and the causal set *C* is unknown

In reality, the set of causal genomic variables *C* is usually unknown. In GWAS, *g*_*ij*_ in (1) was replaced by *h*_*ij*_ that is defined based on all *M* genetic variants:

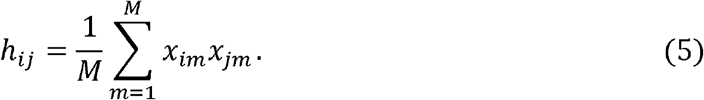

Obviously, *h*_*ij*_ is a noised version of *g*_*ij*_ by including *M* − *K* non-causal genomic variables. Define a *N* × *N* similarity matrix *H* with (*H*)_*ij*_ = *h*_*ij*_. We can derive an estimate based on either REML or PCGC using *H*. Simulations for the REML estimator using GWAS data in early years suggested that these estimates based on *H* provided a practically unbiased estimate for *λ*, which was justified by a theoretical analysis under appropriate assumptions^21^. In what follows, let 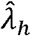 be the PCGC estimator using *h*_*ij*_. In Theorem 2, we will show that 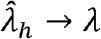 in probability as *M, N* → ∞ in a way that *M*/*N*^2^ → 0. This provides another proof of the work in Jiang et al. (2016)^21^, while their work was based on REML and technically more complicated.

#### Theorem 2.

Let 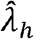 be the PCGC estimator based on *h*_*ij*_ defined in (5). When *M, N* → ∞ in a way that 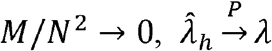.

Now, we consider the scenario when genomic markers are measured with error and the causal set is unknown. Let 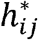 be calculated based on all *M* genomic variables that are measured with error and let 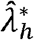 be the PCGC estimator using 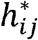. In Theorem 3, we show that the AAF of 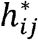 is equal to the average ICC of all genomic variables under appropriate assumptions.

#### Theorem 3.

We assume that *τ*_*m*_ follows the same distribution for causal and non-causal genomic variables. Define 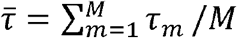 to be the average ICC for all *M* genomic variables. Then, 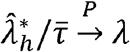 when *M, N* → ∞ in a way that *M*/*N*^2^ → 0.

## 3. Estimating AAF using repeated measurement experiment

When the ICC values *τ*_*m*_ are known for all genomic variables, 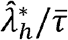 consistently estimates *λ* under appropriate conditions and 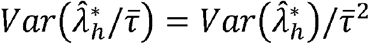. When ICCs are unknown, we need to perform a repeated measurement experiment to derive 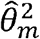, the estimate of the measurement error variance 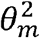. Let 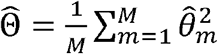 be the average across *M* genomic variables. Since 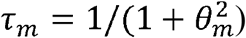, we have the first order Taylor’s expansion:

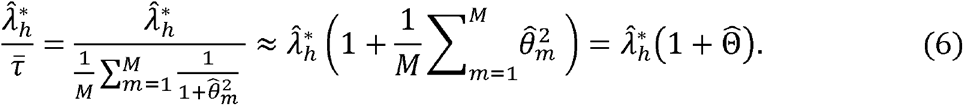

For two independent random variables *z*_*k*_ (*k* = 1,2) with *μ*_*k*_ = *Ez*_*k*_ and 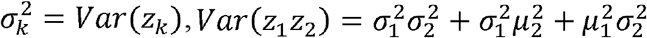. Following this, we have

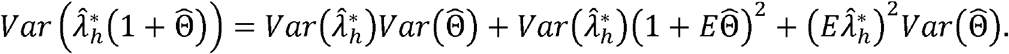

This leads to the following approximation:

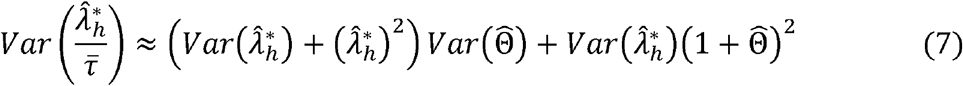

if we replace 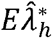 with 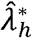 and replace 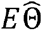 with 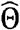. The first term 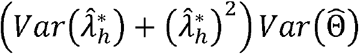 in (7) is the extra variability caused by the uncertainty of the estimated ICC values for *M* genomic variables. Of note, 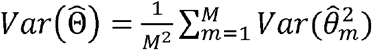 reduces with *M*. Typically, 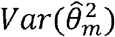 may be quite large for an individual genomic variable due to the limited sample size in the repeated measurement experiment; however, 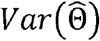 may be small when *M* is large.

## 4. Estimating the correlation of effect sizes between two traits

We consider two studies investigating different traits with *N*_1_ and *N*_2_ subjects, respectively. All subjects are profiled for *M* genomic markers. For subject *i* in study *k* (*k* = 1,2), let *y*_*ki*_ be the trait value and *x*_*kim*_ be the uncontaminated genomic value. All variables are normalized to have mean zero and unit variance. We assume that the same set of *K* genomic markers, denoted as *C*, are causal for the two traits under the following additive models:

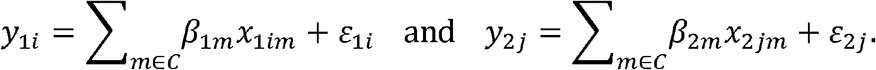

Furthermore, we assume 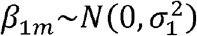 *i. i. d.,*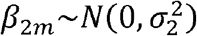*i. i. d*. and *cor*(*β*_1*m*_, *β*_2*m*_) = *ρ* for *m* ∈ *C*. For each trait, 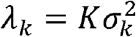 can be estimated using methods in the previous section. Our interest in this section centers on estimating *ρ* when genomic variables are measured with error.

Let 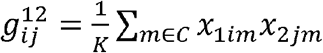 for two subjects *i* and *j* from two studies. We have 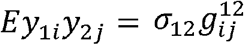, where 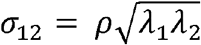. Similarly, we consider regression 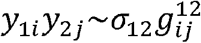 to derive the PCGC estimator

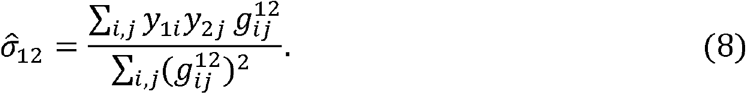

Now, we assume that genomic variables are measured with error, e.g., *x*_*kim*_ is contaminated as 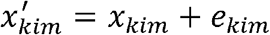 with 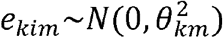. Similarly, we define 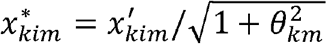 and 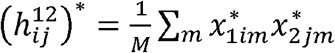 using all *M* genomic variables, and derive the PCGC estimator

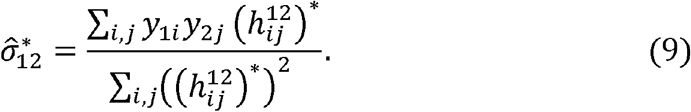

We define 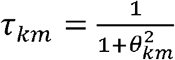 to be the ICC for the *m*^*th*^ genomic marker in study *k* and denote as 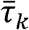 the average ICC across all *M* genomic variables. Let 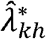 be the PCGC estimator based on all *M* contaminated genomic variables for study *k*. Based on Theorem 3, we have 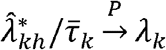. Then, *ρ* can be estimated as 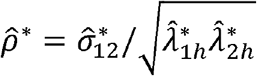.

### Theorem 4.

*For both studies, we assume that the ICC values τ*_*km*_ *are equally distributed between causal and non-causal genomic markers. As N*_1_, *N*_2_, *M* → ∞ *in a way that* 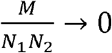,

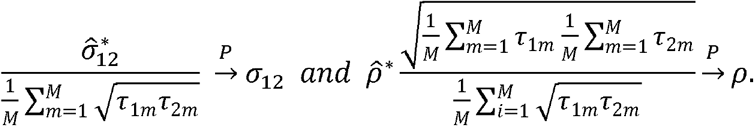

Theorem 4 suggests that the AAF for 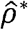 is 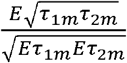. By the Cauchy–Schwarz inequality, AAF ≤ 1 and “=“ holds 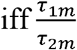 is a constant. In particular, if *τ*_1*m*_ = *τ*_2*m*_ for all markers (a reasonable assumption), AAF=1, i.e., 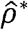 is consistent even when genomic markers are measured with error.

## 5. Simulation studies

In the first set of simulations, we assumed the same measurement error variance (*θ*^2^ = 1 or 0.25, ICC=0.5 or 0.8) across *M* = 10,000 independent variables *x*_*im*_ ∼ *N*(0,1). The fraction of causal variables was set as 100%, 20%, 10% and 5%. Sample size was set at 5000. For each setting, we performed 500 simulations. The results are summarized in Table 1. The naïve estimate without measurement error correction underestimated *λ* with the ICC as the attenuation factor. Our methods provided unbiased estimates after correction.

**Table 1.** Estimates of OFPV *λ* of 10,000 independent predictors measured with a fixed measurement error (ME) standard deviation (*θ*).

| Percentage of<br>causal<br>variables | True $\lambda$ | $\hat{\lambda}$, not corrected for ME | | $\hat{\lambda}$, corrected for ME | |
| --- | --- | --- | --- | --- | --- |
| | | $\theta = 0.5$ | $\theta = 1$ | $\theta = 0.5$ | $\theta = 1$ |
| 100% | 0.1 | 0.082 (0.03) | 0.051 (0.03) | 0.102 (0.04) | 0.103 (0.06) |
|  | 0.5 | 0.400 (0.04) | 0.250 (0.03) | 0.500 (0.05) | 0.500 (0.07) |
|  | 0.9 | 0.719 (0.04) | 0.449 (0.04) | 0.898 (0.06) | 0.898 (0.08) |
| 20% | 0.1 | 0.078 (0.03) | 0.048 (0.03) | 0.097 (0.04) | 0.096 (0.06) |
|  | 0.5 | 0.399 (0.04) | 0.249 (0.03) | 0.499 (0.05) | 0.498 (0.07) |
|  | 0.9 | 0.722 (0.05) | 0.451 (0.04) | 0.902 (0.06) | 0.901 (0.08) |
| 10% | 0.1 | 0.079 (0.03) | 0.050 (0.03) | 0.098 (0.04) | 0.099 (0.06) |
|  | 0.5 | 0.397 (0.04) | 0.248 (0.04) | 0.497 (0.05) | 0.496 (0.07) |
|  | 0.9 | 0.717 (0.05) | 0.447 (0.04) | 0.896 (0.07) | 0.894 (0.08) |
| 5% | 0.1 | 0.079 (0.03) | 0.049 (0.03) | 0.099 (0.04) | 0.099 (0.06) |
|  | 0.5 | 0.400 (0.04) | 0.250 (0.04) | 0.500 (0.06) | 0.500 (0.07) |
|  | 0.9 | 0.722 (0.06) | 0.452 (0.05) | 0.903 (0.08) | 0.904 (0.10) |

In the second set of simulations, we allowed the *M* independent variables to have different measurement error variances 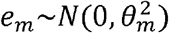, where *θ*_*m*_ were generated from a *Beta*(*a,b*) distribution. We chose (*a,b*) = (3,3) and (5,2) to include symmetric and left skewed distributions, respectively. The average ICC across *M* predictors was 0.793 and 0.664 for the two sets of (*a,b*) parameters, respectively. We repeated the different set-ups where different fractions of predictors were causal. Again, sample size was 5000 and *M* =10,000. These results are summarized in Table 2. Consistent with our theoretical analysis, the attenuation factor was the average ICC across *M* predictors. After correction, we obtained unbiased estimates of *λ* for all simulations.

**Table 2.** Estimates of OFPV *λ* of 10,000 predictors measured with measurement error (ME). The standard deviation of ME follows a beta distribution *θ*∼*Beta*(3,3) or *θ*∼*Beta*(5,2).

| Percentage of<br>causal<br>variables | True $\lambda$ | $\hat{\lambda}$, not corrected for ME | | $\hat{\lambda}$, corrected for ME | |
| --- | --- | --- | --- | --- | --- |
| | | $\text{Beta}(3, 3)$ | $\text{Beta}(5, 2)$ | $\text{Beta}(3, 3)$ | $\text{Beta}(5, 2)$ |
| 100% | 0.1 | 0.078 (0.03) | 0.065 (0.03) | 0.098 (0.04) | 0.098 (0.05) |
|  | 0.5 | 0.396 (0.04) | 0.333 (0.03) | 0.499 (0.05) | 0.501 (0.05) |
|  | 0.9 | 0.715 (0.04) | 0.601 (0.04) | 0.900 (0.05) | 0.904 (0.06) |
| 20% | 0.1 | 0.081 (0.03) | 0.066 (0.03) | 0.102 (0.04) | 0.099 (0.04) |
|  | 0.5 | 0.398 (0.04) | 0.331 (0.03) | 0.501 (0.05) | 0.497 (0.05) |
|  | 0.9 | 0.715 (0.05) | 0.596 (0.04) | 0.900 (0.06) | 0.897 (0.06) |
| 10% | 0.1 | 0.078 (0.03) | 0.065 (0.03) | 0.099 (0.04) | 0.097 (0.04) |
|  | 0.5 | 0.394 (0.04) | 0.33 (0.04) | 0.496 (0.05) | 0.497 (0.06) |
|  | 0.9 | 0.712 (0.06) | 0.597 (0.05) | 0.897 (0.07) | 0.898 (0.07) |
| 5% | 0.1 | 0.083 (0.03) | 0.069 (0.03) | 0.105 (0.04) | 0.105 (0.04) |
|  | 0.5 | 0.397 (0.04) | 0.334 (0.04) | 0.500 (0.05) | 0.503 (0.06) |
|  | 0.9 | 0.710 (0.06) | 0.600 (0.06) | 0.894 (0.08) | 0.902 (0.08) |

In the third set of simulations, we investigated the impact of correlation among predictors. Parameters were the same as the second set of simulations; however, the predictors were simulated to be dependent with correlation coefficient 0.2 or 0.5 for all pairs of predictors. The results are summarized in Table 3. Modest correlation *r* = 0.2 did not impact the estimate of *λ* and we obtained unbiased estimates of *λ* after correction. For the scenario with extensive and strong correlations (*r* = 0.5), the original statistical framework overestimated *λ*. This rarely happens in reality because only few genomic variables are highly correlated and highly correlated variables are typically excluded from analysis. Interestingly, the attenuation factor was still the average of ICC (∼0.667 in current simulations) regardless of the correlation among predictors.

**Table 3.** Estimates of OFPV *λ* of 10,000 dependent predictors measured with measurement error (ME). The standard deviation of ME follows a beta distribution *θ*∼*Beta*(5,2) with average ICC ∼ 0.667. Correlation coefficients between predictors were set at *r* = 0.2 or 0.5.

| Percentage of<br>causal<br>variables | True $\lambda$ | $\hat{\lambda}$, not corrected for ME | | $\hat{\lambda}$, corrected for ME | |
| --- | --- | --- | --- | --- | --- |
| | | $r = 0.2$ | $r = 0.5$ | $r = 0.2$ | $r = 0.5$ |
| 100% | 0.1 | 0.066 (0.029) | 0.085 (0.026) | 0.100 (0.044) | 0.127 (0.039) |
|  | 0.5 | 0.337 (0.034) | 0.429 (0.035) | 0.506 (0.052) | 0.645 (0.053) |
| 20% | 0.1 | 0.066 (0.029) | 0.085 (0.028) | 0.100 (0.043) | 0.127 (0.042) |
|  | 0.5 | 0.334 (0.034) | 0.426 (0.044) | 0.503 (0.051) | 0.641 (0.067) |
| 10% | 0.9 | 0.065 (0.029) | 0.084 (0.028) | 0.098 (0.044) | 0.127 (0.043) |
|  | 0.5 | 0.334 (0.037) | 0.425 (0.05) | 0.503 (0.051) | 0.64 (0.076) |
| 5% | 0.1 | 0.070 (0.031) | 0.087 (0.034) | 0.105 (0.046) | 0.130 (0.051) |
|  | 0.5 | 0.340 (0.062) | 0.427 (0.108) | 0.512 (0.093) | 0.642 (0.163) |

## 6. Real data analysis

The human gut microbiome, also called the second human genome, is the collection of microbes inhabiting the human gut and has been shown to be implicated in many complex diseases in cross-sectional and prospective studies^31^. Estimating the OFPV of human microbiome to the risk of developing a specific disease in prospective studies can help us understand prognostic potential of human microbiome and design future studies appropriately. However, very few prospective microbiome studies are publicly available, and for the ones whose data are available, the sample size is too small to estimate OFPV.

To illustrate the idea of measurement error correction in estimating OFPV, we have used the human gut microbiome data from the American Gut Project (AGP)^32^ to analyze body mass index (BMI) and multiple allergy traits (Table 4). Note that AGP is a cross-sectional study, the estimated *λ* cannot be interpreted as the causal effect of gut microbiome to these traits. Even if the human gut microbiome explains a large fraction of phenotypic variance of, for example, peanut allergy, this may be because individuals with peanut allergy avoid peanut-related food that impacts the composition of gut microbiome communities. Causality needs to be established based on large scale prospective studies. While the causality cannot be established, this type of analyses is important and informative for microbiome studies^33^.

**Table 4.**
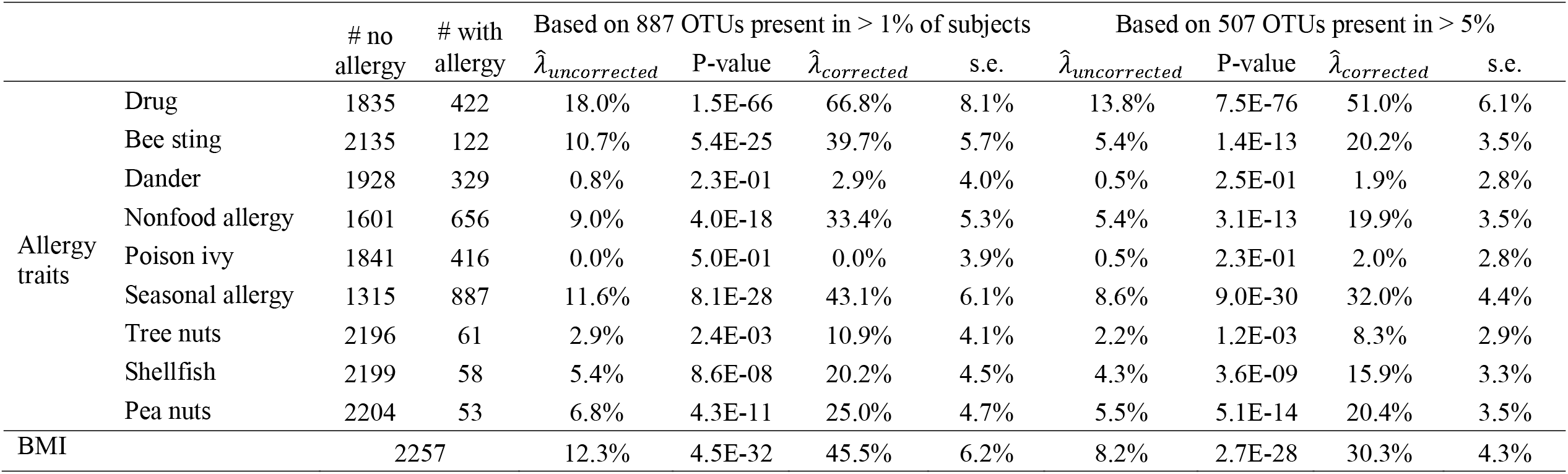
Estimated phenotypic variance attributed to gut microbiome in American Gut Project.

The AGP data, including the operational taxonomy units (OTU) table and clinical variables, were downloaded from the Github website (https://github.com/biocore/American-Gut/tree/master/data/AG). We calculated the relative abundance (RA) table based on the OTU table. In total, 2257 subjects were available for analysis after filtering (sample exclusion: age missing or ≤3, history of using antibiotics in the past month, patients with type 2 diabetes or inflammatory bowel disease, or the number of reads < 1000 after quality control).

We estimated measurement error variances and ICCs for taxa using data with the repeated measurement data separated by approximately six months from the Human Microbiome Project^34^ (HMP). For 887 taxa present in at least 5% of HMP subjects and at least 1% in AGP subjects, we fitted a random effect model using an R package *lme4*^*35*^ to estimate measurement error variances and calculated ICCs. The average ICC for those taxa was found to be 0.27.

We performed two sets of analyses. The first set included 887 OTUs present in at least 1% of AGP subjects while the second set included 507 OTUs present in at least 5% of HMP subjects (average ICC=27%). Results are summarized in Table 4. Overall, human gut microbiome was significantly associated with BMI and all allergy traits except for dander and poison ivy allergy, consistent with our previous beta-diversity analyses^36^. ICCs were relatively low in general for microbiome primarily due to temporal instability; ignoring measurement error may severely underestimate the overall contribution of microbiome to the trait. For example, correcting for measurement error increased the estimate for seasonally allergy from 11.6% to 43.1% using 887 taxa, suggesting that a very large fraction of seasonally allergy is related to gut microbiome. **In our analyses, gut microbiome explained 12.3% of BMI variance without correction for measurement error and 45.5% after correction. A recent paper estimated that 25% of BMI variance was attributed to gut microbiome**^**33**^ **without considering correcting for measurement error. This uncorrected estimate was higher than our estimate possibly because of different covariates adjusted in the model**.

Finally, we point out the estimates would be higher when we include more predictors at the price of increased standard error. For example, for seasonal allergy, the measurement error-corrected estimate 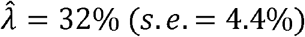 using 507 OTUs present in at least 5% subjects and 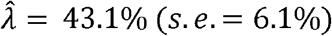 using 887 OTUs present in at least 1% subjects.

## 7. Discussion

Prospective studies examining the association between high-dimensional genomic variables and different diseases are essential for understanding disease etiology and for building risk prediction models. Estimating the OFPV of a disease/trait attributed to all genomic variables is a crucial step to understand the upper limit of the potential of genomic variables for predicting a disease and design future projects. We have investigated the impact of measurement error on estimating OFPV. Our analysis shows that AAF equals to the average ICC across all causal genomic variables. Under the assumption ICCs are distributed similarly between causal and non-causal variables, AAF equals to the average ICC of all genomic variables.

When ICCs are known in advance, we can consistently estimate OFPV and adjust the variance of the estimator. When ICCs are unknown, we have to estimate ICCs using pilot studies with repeated measurements. The sample size of a pilot study is often limited; thus, the uncertainty of the estimated measurement error variances is large for individual variables. However, since AAF is the average ICC across all genomic markers, the extra variance due to the uncertainty of ICC estimate is small if the number of genomic variables is large.

In GWAS, measurement error comes from both genotyping and imputation. For common variants, since only well-genotyped and well-imputed variants are used, measurement error is expected to have a relatively small impact on heritability analysis. As large-scale sequencing studies are developed, heritability analyses are performed for rare variants derived from sequencing or imputation. In this case, the impact of measurement error of heritability analysis must be evaluated. For other genomic variables (e.g., microbiome), measurement error would have relatively large impact on estimating OFPV and must be accounted for.

Further, we have derived the AAF for estimating the correlation of effect sizes of *M* variables on two traits (also called co-heritability in genetics). We have considered two independent sets of subjects profiled for the same set of genomic markers. A similar analysis can be applied to a single set of subjects with both traits and the conclusion remains the same. As we show, when 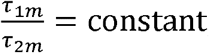, i.e., the ratios of ICCs between two studies are the same across all variables, AAF equals to one. In reality, we expect the same ICC for a genomic variable in both studies; thus, we do not need to adjust for measurement error for estimating this parameter.

There are several limitations in the current analysis. First, we assume that genomic variables are independent, which may not hold for most of genomic studies. Our numerical study shows that modest correlation does not seem to bias the estimate and the conclusion of measurement error correction is still valid. Thus, for real data analysis, we would suggest performing correlation-based pruning so that no extensively highly correlated genomic variables are included for the analysis. Future theoretical analysis is required for any correlation structure among genomic variables. Second, our analysis was based on the PCGC estimator, a type of moment estimator. This estimator is less efficient compared to the REML estimator; it would be interesting to extend current analysis to the REML framework. Third, the current analysis is for a quantitative trait, and it will be interesting to extend it to survival traits in cancer genomics and case-control studies by accounting for ascertainment^7,11^. Moreover, we did not consider the measurement error in fixed effects. Finally, we assume that measurement error is additive and independent, which needs to be investigated empirically. Cui and his colleagues^19^ assume a general covariance structure for measurement error; however, it is impossible to apply their approach to a study with a large number of genomic variables. Future research is warranted in this direction.

## Appendix A.

### Notations and lemmas

For convenience, we define

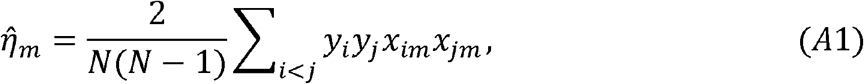

the numerator of the PCGC estimator based on a single genomic variable. One can verify that 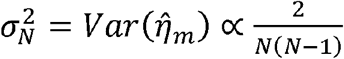. As 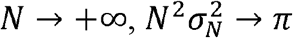, a finite constant. Some algebra leads to

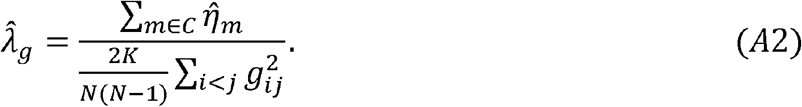

We will show that the denominator 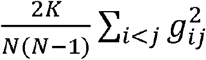 converges to 1 when *g*_*ij*_ is calculated based on *K* independent predictors. Thus, 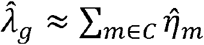 and the representation (A2) greatly simplifies the presentation for our technical analysis.

#### Lemma 1.

Assume random variables (RVs) *X*_1_, ⋯, *X*_*N*_ *i*.*i*.*d*. with *EX*_*i*_ = 0, *Var X*_*i*_ = 1 and 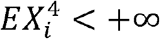. Then 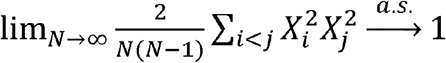.

Proof. Some algebra leads to 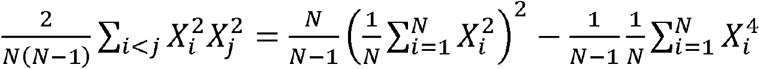. As 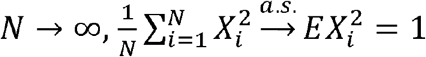 and 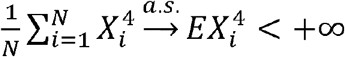. This completes the proof.

#### Lemma 2.

Assume RVs *X*_*im*_ (*i* =1, ⋯, *N*; *m* =1, ⋯, *M*) *i*.*i*.*d*. with *EX*_*im*_ = 0, *Var X*_*im*_ = 1 and 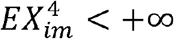. Then 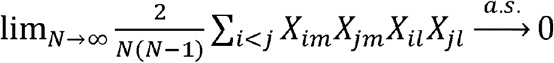.

Proof. Note that 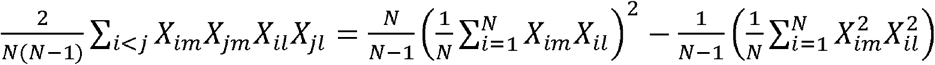. As *N* → ∞, this converges to 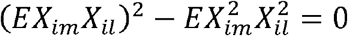 because of independence.

#### Lemma 3.

Assume RVs *X*_*im*_ (*i* =1, ⋯, *N*; *m* =1, ⋯, *M*) *i*.*i*.*d*. with *EX*_*im*_ = 0 and *Var X*_*im*_ = 1 and 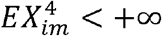. Let 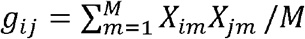. Then,

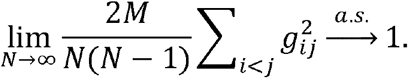

Proof. One can verify that the above equation equals to

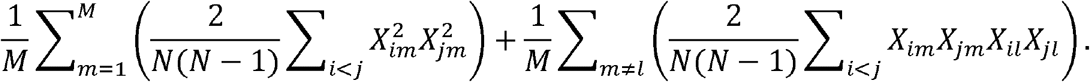

As *N* → ∞, the first term 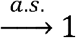 1 by Lemma 1 and the second term 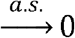 0 be Lemma 2. This completes the proof.

#### Lemma 4.

Suppose independent 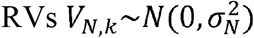 with 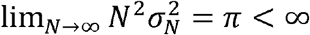. Let 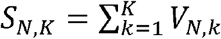. Then, 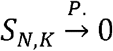 as when *N, K* → ∞ in a way that *K*/*N*^2^ → 0.

Proof. When *N, K* → ∞ in a way that *K*/*N*^2^ → 0, 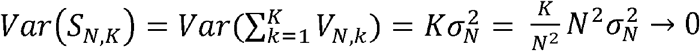 because 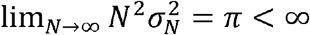. This implies 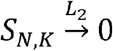 and thus 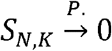.

#### Lemma 5.

Let *λ* > 0 be a fixed constant. Given *K*, let *V*_1_ ⋯, *V*_*K*_ be independent RVs with 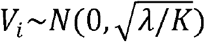. Given *V*_*i*_, define *W*_*i*_ to be a RV with 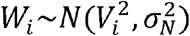 and 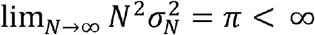. Let *U*_1_ ⋯, *U*_*K*_ be *i*.*i*.*d*. RVs with 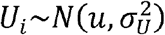 and 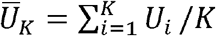. When *N, K* → ∞ in a way that 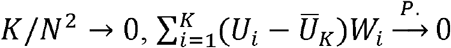.

Proof. Let 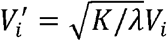, then 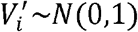. Since 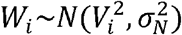, we assume 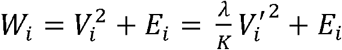, where 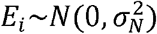. Thus,

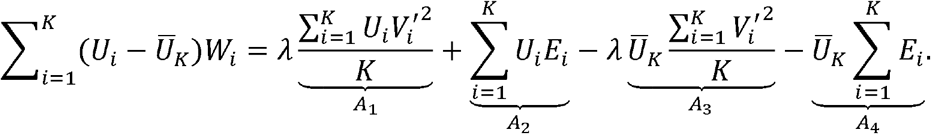

By the law of large numbers, as 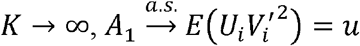 and 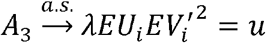; thus, 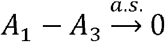. Moreover, because 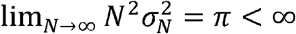 and 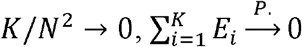 (i.e., 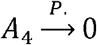) and *A*_2_ → 0 by Lemma 4. This completes the proof.

## Appendix B.

### Proof of Theorem 1

Proof. By definition,

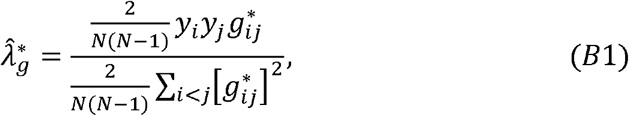

where 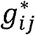 defined in (4) is based on error-prone predictors. Combining (3) and (4), we have

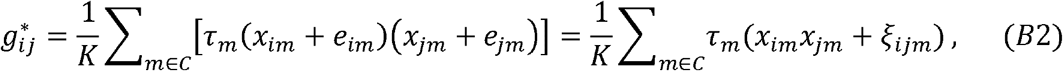

where *ξ*_*ijm*_ = *x*_*im*_*e*_*jm*_ + *e*_*im*_*x*_*jm*_ + *e*_*im*_*e*_*jm*_. Plugging (*B2*) into (2) and noticing 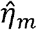 defined in *(A1)*, we found that 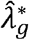 can be rewritten as:

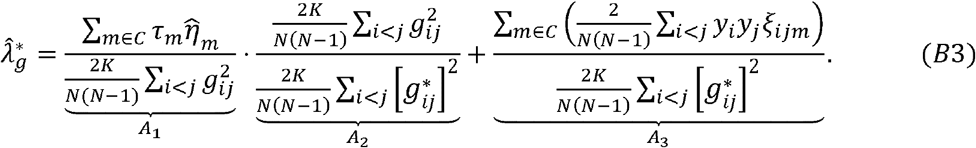

By Lemma 3, the numerator and denominator of *A*_2_ converge to 1 and thus *A*_2_ → 1 as *N* → 0. We next show that 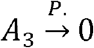. Since the denominator of 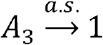 by Lemma 3, it remains to prove that the numerator converges to zero in probability. It is easy to verify that *cov*(*y*_*i*_*y*_*j*_, *ξ*_*ijm*_) = 0, thus 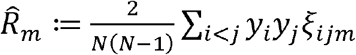 is the PCGC estimator for zero using data *ξ*_*ijm*_; thus, 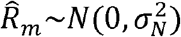 asymptotically with 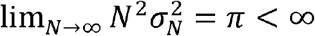. Thus, the numerator of *A*_3_ becomes 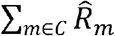 and converges to zero in probability by Lemma 4.

Finally, we prove that 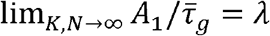. Note that *A*_1_ can be rewritten as

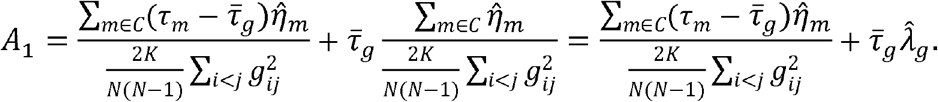

The second equation holds because of (A2). Note that 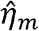 is the PCGC estimator for 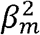. This suggests that 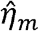 can be written as 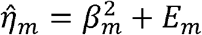, where *β*_*m*_ ∼ *N*(0, *λ*/*K*)is the effect size for the variable in the random effect model and 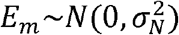. By Lemma 5, 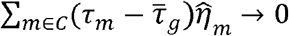 (and thus the first term of *A*_1_ → 0) as *N, K* → ∞ in a way that *K*/*N*^2^ → 0. Since it is established that 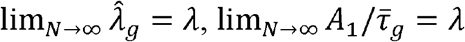. This completes the proof.

## Appendix C.

### Proof Theorem 2

Proof. By definition,

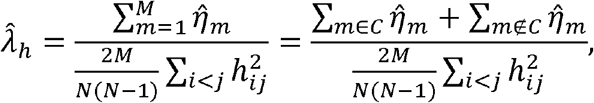

which can be further written as

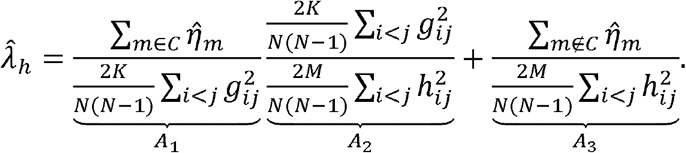

It is established that 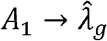. Also, *A*_2_ → 1 because both the numerator and the denominator converge to 1 as *N* → + ∞. Thus, *A*_1_*A*_2_ → *λ* as *N* → + ∞. Moreover, by Lemma 4, we have 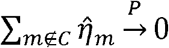 (as thus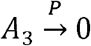) as *M, N* → + ∞ in a way that *M*/*N*^2^ → 0. This completes the proof.

## Appendix D.

### Proof of Theorem 3

Proof. Let *τ*_0_ = *Eτ*_*m*_. Note that,

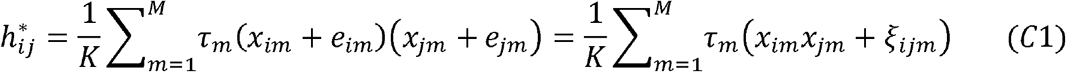

where *ξ*_*ijm*_ = *x*_*im*_*e*_*jm*_ + *e*_*im*_*x*_*jm*_ + *e*_*im*_*e*_*jm*_. Then, 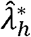 can be rewritten as:

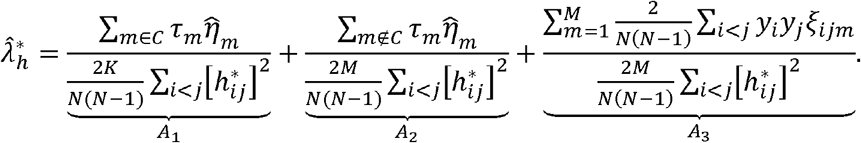

Note that 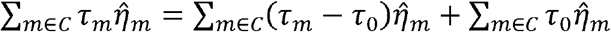. Using a similar argument as in Theorem 1, we have *A*_1_ → *τ*_0_*λ*. Moreover, because 0 ≤ *τ*_*m*_ ≤ 1, we have 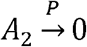 based on Lemma 4. Finally, using the same argument for *A*_3_ in Theorem 1, 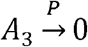 when *M, N* → + ∞ in a way that *M*/*N*^2^ → 0. This completes the proof.

## Appendix E.

### Proof of Theorem 4

Proof. Using a similar argument as in Lemma 3, we can prove that the denominator of 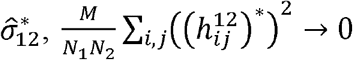, as *N*_1_, *N*_2_ → + ∞. So, we ignore the denominator for simplicity.

Let 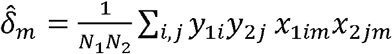. Let 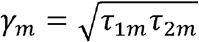 and *γ*_0_ = *Eγ*_*m*_. Then,

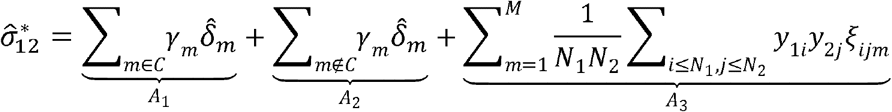

Note that 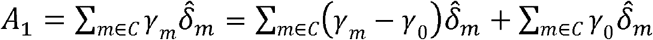. Using a similar argument as in Theorem 1, we have 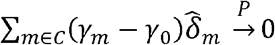 and thus 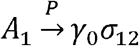 as *N*_1_, *N*_2_, *M* → ∞ in a way that *M*/*N*_1_*N*_2_ → 0. Moreover, because 0 ≤ *γ*_*m*_ ≤ 1, we have 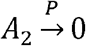based on Lemma 4. Finally, using the same argument for *A*_3_ in Theorem 1, 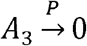 as *N*_1_,*N*_2_,*M* → ∞ in a way that *M*/*N*_1_*N*_2_ → 0. This completes the proof.

## Acknowledgements

This work utilized the computational resources of the NIH HPC Biowulf cluster. (http://hpc.nih.gov). The authors are supported by the NIH Intramural Research Program.

